# Comprehensive Phenotyping of Cutaneous Afferents Reveals Rapid-Onset Alterations in Nociceptor Response Properties Following Spinal Cord Injury

**DOI:** 10.1101/2021.06.15.448577

**Authors:** Olivia C. Eller, Rena N. Stair, Christopher Neal, Peter S. N. Rowe, Jennifer Nelson-Brantley, Erin E. Young, Kyle M. Baumbauer

**Author notes:** **Corresponding Author:** Dr. Kyle M. Baumbauer, Department of Anatomy and Cell Biology, University of Kansas Medical Center, 3901, Rainbow Blvd, MS 3051, Kansas City, KS 66160.

## Abstract

Spinal cord injury (SCI) is a complex syndrome that has profound effects on patient well-being, including the development of medically-resistant chronic pain. The mechanisms underlying SCI pain have been the subject of thorough investigation but remain poorly understood. While the majority of the research has focused on changes occurring within and surrounding the site of injury in the spinal cord, there is now a consensus that alterations within the peripheral nervous system, namely sensitization of nociceptors, contribute to the development and maintenance of chronic SCI pain. Here we demonstrate that thoracic spinal contusion injury results in the emergence of autotomy and spasticity, both indicators of spontaneous pain, in areas below the level of the injury within 24 hr of SCI. These behaviors were associated with hindpaw edema and elevated cutaneous calcitonin gene-related peptide (CGRP) concentration. Electrophysiological recordings using an *ex vivo* skin/nerve/DRG/spinal cord preparation demonstrated that SCI increased mechanical and thermal sensitivity, as well as the incidence of spontaneous activity (SA) and afterdischarge (AD), in below-level C-fiber nociceptors 24 hr following injury. Interestingly, the distribution of nociceptors that exhibit SA and AD are not identical, and the development of SA was observed more frequently in nociceptors with low thermal thresholds, while AD was found more frequently in nociceptors with high thermal thresholds. These results demonstrate that SCI causes the rapid-onset of peripheral inflammation-like processes that sensitize nociceptors, which may contribute to the early emergence and persistence of chronic SCI pain.

## 1. Introduction

Spinal cord injury (SCI) can have devastating consequences that are most commonly associated with loss of motor function. Individuals with SCI experience a number of additional complications, with the majority of patients experiencing chronic pain that is often severe. [13; 16; 21; 29; 39; 52; 62; 63; 71]. This is significant because ongoing pain may impair recovery for those undergoing rehabilitation [5-7; 34; 37]. Unfortunately, current pain management relies on opioids, despite research demonstrating that these drugs are unsuccessful at treating chronic pain and can even exacerbate functional loss associated with injury (e.g., bowel motility), potentiate the development of pain, and undermine locomotor recovery. [1; 3; 14; 30; 38; 83]. Consequently, there is a need to better understand the mechanisms underlying SCI pain to improve treatment options and patient quality of life.

Pain following SCI is identified regionally by the relationship of pain experience to the site of injury. At-level pain is reported in dermatomes and myotomes that are innervated by sensory neurons originating from the injury site. Below-level pain, by comparison, originates from tissue below the level of injury [35; 74], and is unique because it occurs despite disrupted connectivity in the ascending neural pathways and in the absence of normal somatosensation [70]. This type of pain occurs in the majority of SCI pain patients [15], and is characterized as a spontaneous, intense burning or stabbing sensation that is refractory to medical intervention [15].

While the mechanisms underlying below-level pain are complex, there is now evidence demonstrating that nociceptors contribute to the development and maintenance of SCI-induced pain [8; 9; 17; 89]. Furthermore, while the majority of SCI pain isn’t reported for weeks to months after injury, research suggests that increases in nociceptor excitability, such as the emergence of spontaneous activity (SA), may occur within days following SCI. SA is known to underly spontaneous pain [9; 25; 26; 54; 56; 61; 79; 89], however, there has been little research examining whether the emergence of SA is also associated with rapid development of other types of aberrant nociceptor activity, such as afterdischarge (AD), or whether SCI alters response properties in nociceptors with specific functional response properties. We show that spontaneous below-level pain, hindpaw edema, and increased cutaneous calcitonin gene related peptide (CGRP) appear 1-7 days following SCI, suggesting that SCI produces peripheral inflammation during the acute phase of injury. To determine whether these inflammation-related alterations were also associated with changes in nociceptor function, we performed *ex vivo* electrophysiological recordings from cutaneous nociceptors and found that SCI increases mechanical and thermal sensitivity, as well as the incidence of SA and AD in unmyelinated nociceptors. Further, SA and AD were associated with thermal, but not mechanical, sensitivity. This work is an important step in establishing the mechanisms underlying SCI pain by further supporting the role of the peripheral nervous system in the development of pathological pain following SCI.

## 2. Methods

### 2.1. Animals

Experiments were conducted using 8-12 week-old (20-30 g) female C57BL/6J mice (Jackson Laboratories, Bar Harbor, ME) that were group housed (4-5 mice/cage) and maintained in a temperature-controlled environment on a 12:12 light:dark cycle with free access to food and water. All studies were approved by the UConn Health and University of Kansas Medical Center Institutional Animal Care and Use Committee and treated in accordance with published NIH standards.

### 2.2. Spinal contusion injury

Mice were randomly assigned to SCI and naïve conditions and SCI mice were anesthetized with isoflurane (2%). A sham condition was not included because prior work has shown that laminectomy alone is sufficient to produce nociceptor sensitization and pain in mice [56; 57]. A rostro-caudal incision was made to expose the thoracic level vertebrae and surrounding musculature. Tissue was carefully removed from around the vertebral column, and a laminectomy was performed at the level of T8-T11 using fine surgical scissors and forceps (Fine Science Tools; Foster City, CA). The spinal column was then secured into position on the impactor using modified forceps (Fine Science Tools; Foster City, CA) clamped to vertebral tissue rostral and caudal of T8 and T11, respectively. Dorsal spinal contusions were produced at the T9 spinal level using the Infinite Horizons IH-0400 Spinal Cord Impactor (Precision Systems and Instrumentation; Lexington, KY) equipped with a standard mouse impact tip (1.3 mm diameter) at a severity of 65 kDynes and a 1 s dwell time. This level of injury is sufficient to produce hindlimb paralysis that results in a moderate level of recovery of locomotor function over time. The T9 spinal level was chosen to leave the lumbar spinal cord and dorsal root projections intact. Following the contusion, the wound was closed with 7-0 vicryl sutures, mice were injected with 2 mL of 0.9% saline and 5 mg/kg of gentamicin sulfate and were placed on warm heating pads until regaining consciousness. After displaying ambulatory behavior, mice were individually housed in cages with enrichment materials (plastic huts, burrowing material, etc.) and food, water, and supplemental gel diet (ClearH_2_O, Westbrook, ME) placed in specially designed containers on the cage floor. Cages were placed on warm heating pads such that half of the cage remained off of the heating pad in quiet rooms away from foot traffic and were isolated from other animals for the 7 day duration of experiments. Mouse bladders were manually voided twice daily and supplemental injections of 0.9% saline and antibiotic were administered for 5 days as described in [4]. Bladder voiding was discontinued if reflexive bladder emptying recovered.

### 2.3 Micro computed tomography (µCT) and spinal cord contrast imaging

SCI and naïve mice were anesthetized with a lethal exposure to isoflurane and intracardially perfused with ice cold 0.9% saline followed by 4% paraformaldehyde solution in phosphate buffered saline (PFAPBS). Whole spinal cords were carefully removed, post-fixed for 24 hours in PFAPBS at 4°C, and then suspended in a solution containing 260 mg/mL iohexol (Omnipaque^TM^; GE Healthcare, Marlborough, MA) in 70% ethanol at 4°C for 7 days. Spinal cords were then wrapped and heat-sealed in cling film to prevent dehydration and stacked in a µCT sample container (12 mm diameter x 75 mm height) for batch analysis. Samples were scanned using a Scanco µCT40 imaging system supported by a Hewlett Packard Integrity itanium Server rx2660 with dual core itanium processors 32 GB memory and Open VMS operating system (Scanco Medical USA, Inc – Southeastern, PA 19399). The cone-beam µCT40 system has a peak energy input range of 30 to 70kVp, a maximum scan/sample size 36.9 x 80 mm with a 6 µm resolution capability. A specific *batch control file* for spinal cord contrast analysis was used with the following specifications: **1.** Energy Intensity 45 kVp, 177 µA and 8 W; **2.** A FOV/Diameter of 12 mm with a voxel (VOX) resolution size of 6 µM; **3.** Integration time 300 mS and, **4.** Data Averaging = 7. Following raw data acquisition and computer reconstruction the 6 µM *.DCIM output files were contoured and defined using the “Scanco” software morph or integration functions. For 3D image and quantitative analysis, a script file was set at Gauss Sigma = 4 and Gauss Support = 8 for all thresholds. Scanning thresholds for the spinal anatomic structures were set at: **1**. *Dorsal/lateral/ventral horn definition* = Lower Threshold = 350 permille and Upper Threshold 434 permille; **2.** *Posterior/anterior/lateral funiculus definition* = Lower Threshold = 79 permille and Upper Threshold = 350 permille; and **3.** Central canal/anterior fissure Upper threshold < 79. The Scanco software “morpho” script command file was then used to generate a quantitative 3 Dimensional Seg.Aim file of the region of interest (ROI). Further descriptions are provided in our previous publications [22; 24; 49; 64; 65; 75; 91-93].

Processing of *.DCIM files for image reconstruction and analysis was performed in 3D Slicer (http://www.slicer.org). Segments were created using the Segment Editor module. An Otsu automatic threshold was applied to separate the complete cord volume from the background, and the upper and lower threshold limits were adjusted manually. The largest, single volume was kept using the “islands” sub-menu command, which resulted in the removal of all the smaller, non-cord volumes. Any remaining artifacts were manually detached from cord using the “eraser” tool, and the primary cord volume was kept, repeating the previously used islands command. These steps were repeated on a second segment, which was created for the grey matter. This pair of steps for creating volumes for the grey and white matter for a given cord was repeated for each master volume segment until the complete length of the imaged cord was segmented. Measurements of gray and white sparing, total volume, surface area, roundness, flatness, and elongation were calculated based on the volume labelmap in the “Segment Statistics” module.

### 2.4. Ex vivo preparation and electrophysiological recordings

The *ex vivo* skin/nerve/DRG/spinal cord preparation (depicted in **Figure 1**) is similar to what has been described elsewhere [44; 46; 53; 84]. Briefly, mice were anesthetized with a 90/10 mg/kg mixture of ketamine and xylazine (i.m.) and were transcardially perfused with ice cold oxygenated (95% O2–5% CO2) artificial cerebrospinal fluid (aCSF; in mM: 1.9 KCl, 1.2 KH2PO4, 1.3 MgSO4, 2.4 CaCl2, 26.0 NaHCO3, 10.0 D-glucose, and 127.0 NaCl). The spinal cord, saphenous nerve, DRG, and hindpaw skin were dissected in continuity and transferred to a recording chamber also containing oxygenated aCSF. The skin was then placed on an elevated metal platform with the epidermis exposed to air for mechanical and thermal stimulation. The dermis was continually perfused with bath aCSF that was maintained at 31°C. Electrophysiological recordings were performed by impaling individual sensory neuron somata with sharp quartz microelectrodes containing 1 M potassium acetate. Orthograde electrical search stimuli were administered through a suction electrode placed on the saphenous nerve to locate sensory neuron somata innervating the skin. Receptive fields (RF) were localized on the skin using mechanical (e.g., paintbrush) or thermal stimuli (∼51°C or ∼0°C 0.9% saline). Response characteristics of DRG neurons were determined by applying digitally controlled mechanical and thermal stimuli. The mechanical stimulator consisted of a tension/length controller (Aurora Scientific, Aurora, ON, Canada) attached to a 1-mm diameter plastic disk. Computer controlled 5-s square waves of 1, 5, 10, 25, 50, and 100 mN were applied to the cells RF. After mechanical stimulation, a controlled thermal stimulus was applied using a 3-2 mm contact area Peltier element (Yale University Machine Shop, New Haven, CT, USA). The temperature stimulus consisted of a 12 s heat ramp from 31°C to 52°C, followed by a 5-s holding phase, after which the temperature was ramped back down to 31°C over a 12 s period. A 30 s resting period was inserted between stimulus presentations. In some instances, fibers could not be characterized by computer-controlled mechanical and/or thermal stimulation but were able to be phenotyped using von Frey and/or saline stimuli, respectively. These cells were not included in threshold determination. All elicited responses were recorded digitally for offline analysis (Spike 2 software, Cambridge Electronic Design, Cambridge, UK).

**Figure 1.**
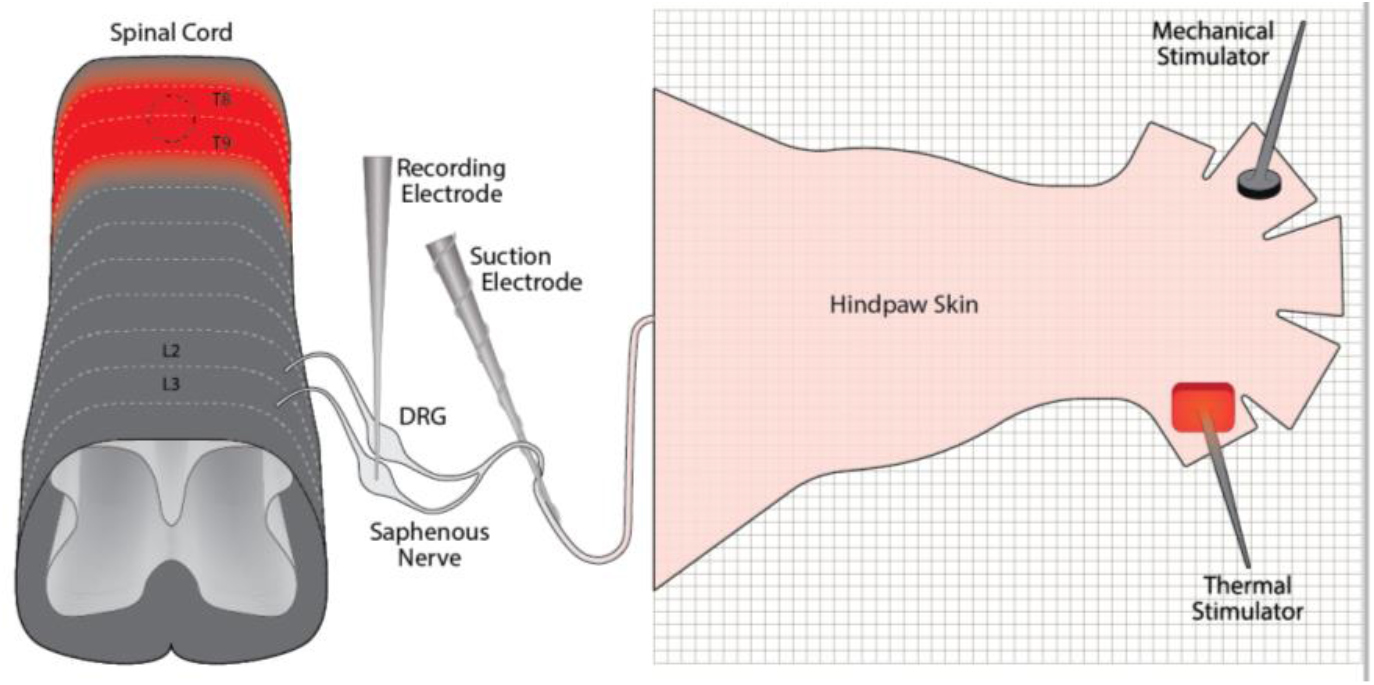
Schematic depicting *ex vivo* skin/nerve/DRG/spinal cord preparation. Spinal contusion injury was performed at T8-T9. Spinal cord tissue rostral to the lesion site and caudal of L4 was dissected, leaving L2 and L3 DRG and the saphenous nerve in continuity with the hairy skin of the hindpaw. Sharp electrode recordings were performed by impaling neuron cell bodies contained within L2 and L3 DRG while heat, cold, and mechanical stimuli were delivered within the cutaneous receptive field of the neuron.

Peripheral conduction velocity was then calculated from spike latency and the distance between stimulation and recording electrodes (measured directly along the saphenous nerve). A-fibers were identified as sensory neurons with conduction velocities > 1.2 m/s, while C-fibers were identified as sensory neurons with conduction velocities of <1.2 m/s [47; 48]. C-fibers were classified as follows: (1) those that responded to mechanical and heat stimuli (CMH); (2) cells that responded to mechanical, heat and cold stimulation (CMHC); (3) those that responded only to mechanical stimulation of the skin (CM); (4) those that responded to mechanical and cooling stimuli (but not heating) (CMC); (5) those that were cold and mechanically insensitive, but heat sensitive (CH); and (6) those that were heat and mechanically insensitive but responded to cooling of the skin (CC).

### 2.5. Enzyme-Linked Immunosorbent Assays (ELISA)

Mice were anesthetized with a lethal exposure to isoflurane and intracardially perfused with ice cold 0.9% saline prior to the dissection and collection of hindpaw skin. Hindpaw hairy skin was homogenized in ice-cold RIPA buffer/protease inhibitor cocktail and centrifuged for 20 min (4^◦^C; 18,000 rcf). Each sample’s total protein concentration was determined using Pierce BCA Protein Assay Kit (Thermo Fisher Scientific; Waltham, MA). ELISAs for CGRP (MyBioSource; San Diego, CA), substance P (SP; Enzo; Farmingdale, NY), and nerve growth factor β (NGFβ; Boster Bio; Pleasanton, CA) were run according to manufacturer’s instructions. All samples were run in duplicate and absorbance ratios were read at 450 nm. Protein concentration was determined by comparison to linearized protein concentration standards, and analyses were conducted using the mean concentration for the duplicate wells for each sample.

### 2.6. Statistical Analysis

Data were analyzed using Student’s t-test, and one-way or mixed designs univariate analysis of variance (ANOVA). *Post hoc* testing was performed using Tukey honestly significant difference (HSD) t-tests where appropriate. Afferent population data were analyzed using Chi square (χ^2^), and linear regression was used to determine relationships between nociceptor response thresholds and alterations in spontaneous firing, as well as SA and AD. Significance was determined using *p* values ≤ .05.

## 3. Results

### 3.1. Site of SCI does not include lumbar spinal locations of saphenous nerve termination

Location and characterization of thoracic contusion injury was performed using 3D reconstructions of μCT images (n=3/condition). The epicenter of the lesion was determined by identifying where the defining characteristics of the dorsal and ventral horns were absent. Once the lesion epicenter was identified, total length of the lesion was calculated by measuring the linear distance between healthy rostral and caudal segments. Because there was variation in the total lesion site length, we divided lesion sites into 5 subsections (20% of total length) to normalize where measurements were taken for comparisons between conditions. The 0-20% bin was the most rostral segment, the 40-60% bin contained the lesion epicenter, and the 80-100% was the most caudal segment (**Figures 2A-C**). We then measured white (WMS) and gray matter sparing (GMS) and found that WMS was significantly reduced while GMS was increased around the lesion epicenter 7 days following SCI, all Fs >7.10, *p* <.05 (**Figures 2D, E**). No significant relationships were detected in any of the other measures taken. Importantly, we were able to verify that the lesion did not extend into L2-L3 spinal segments [36], where saphenous nerve endings terminate.

**Figure 2.**
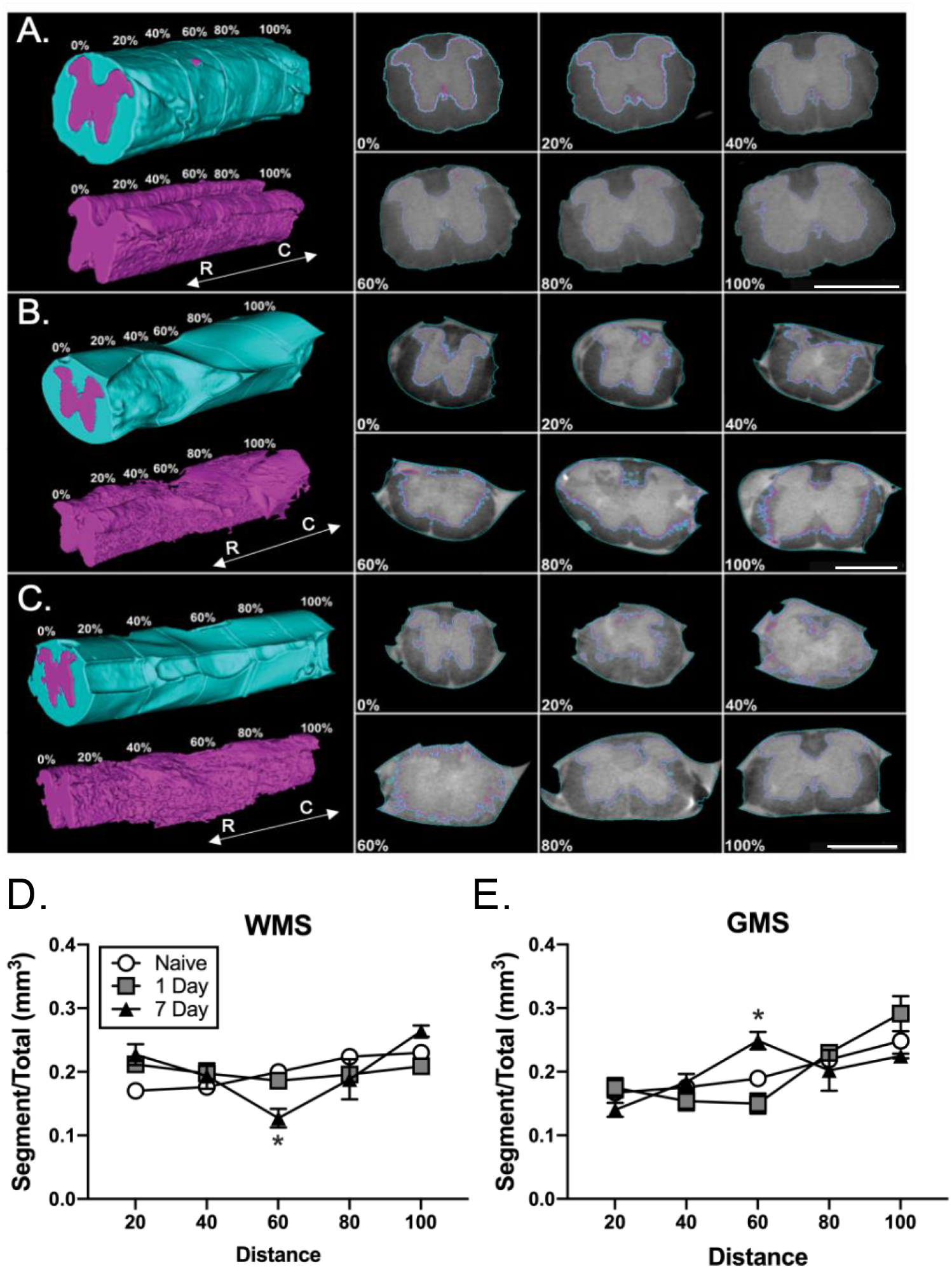
Volumetric reconstruction (μCT) of naïve (**A**), SCI 1 day (**B**), and SCI 7 day (**C**) spinal cords depicting total longitudinal lesion volume, gray mater longitudinal volume, and cross-sections taken at increments of 20% of the total cord length. 0% represents the most rostral healthy margin, 100 % represents the first rostral margin of healthy tissue, and 40-60% captures the lesion epicenter. Gray matter outlined in purple, scale bar = 1 mm. Quantification of white matter sparing (WMS; **D**) and gray matter sparing (GMS; **E**) for each segment of the lesion.

### 3.2. SCI increases the occurrence of spontaneous pain behaviors in injured mice

SCI is often accompanied by the development of spontaneous pain, indicated by the presence of autotomy and muscle spasticity [10; 11; 27; 28; 32; 42; 45; 90]. Therefore, we assessed the presence of both behaviors in a cohort of 9 SCI mice for 7 days following SCI. We found that 5/9 SCI mice exhibited signs of autotomy during this time period, with wounding to the lower abdomen, hindpaws, and/or tail. We also observed hindlimb spasticity in 3/9 mice, all of which also exhibited autotomy (**Figure 3**), further suggesting an overlap in underlying mechanism for these two behavioral outcomes. Neither autotomy nor spasticity were observed in naïve mice (n=8).

**Figure 3.**
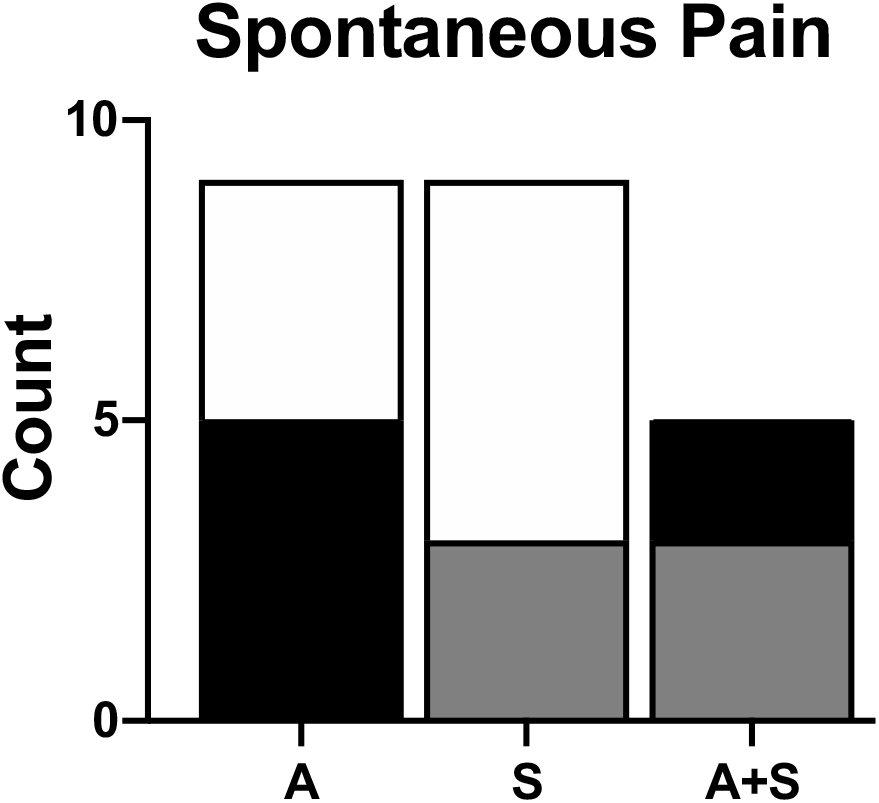
Quantification of autotomy and spasticity for 7 days following SCI. Autotomy and spasticity were characterized as being present or absent in each of 9 SCI mice. The graph depicts the total number of mice that exhibited autotomy (A; black) and spasticity (S; gray) as a proportion of the total number of mice (white). The final bar depicts the number of mice in the autotomy group (black) that also exhibited spasticity (A+S; gray).

Inflammation is a significant driver of both central and peripheral pain states [18; 33; 50; 59; 77; 78]. In the periphery, inflammation is often observed in the form of edema. To determine whether hindpaw autotomy was associated with inflammatory processes that developed during the acute phase of SCI, we measured paw thickness in a separate cohort of 10 naïve and 15 SCI mice during the first 7 days of recovery following SCI. We found that hindpaw diameter increased significantly following SCI relative to naïve mice. Latency (days) to reach peak hindpaw thickness did not differ between mice, however average paw thickness was significantly greater in SCI (2.20 ± 0.09 mm) relative to naïve (1.85 ± 0.09 mm) mice, F(1,24)=7.34, *p*<.05 (**Figure 4A**).

**Figure 4.**
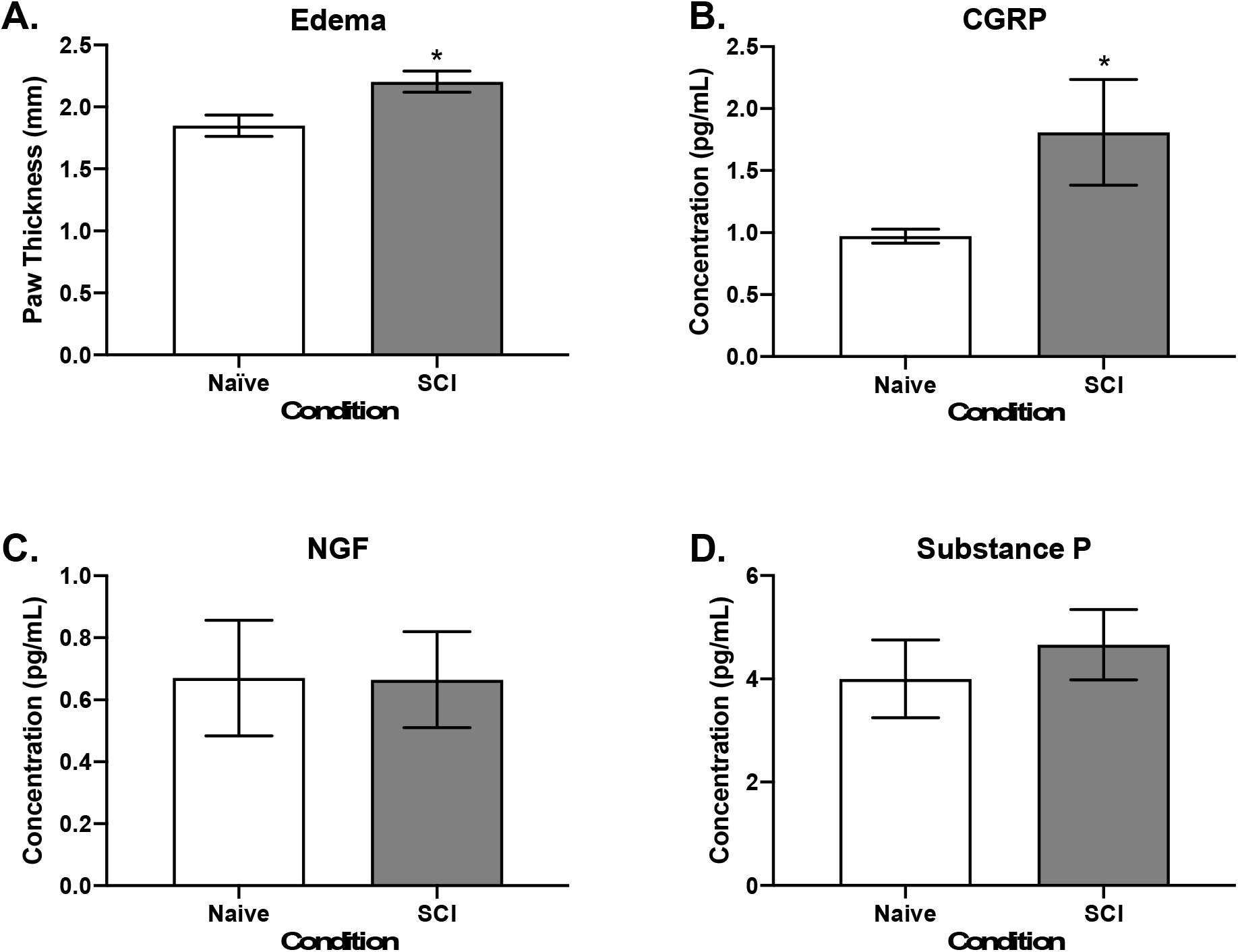
Analysis of hindpaw edema and concentrations of cutaneous CGRP, NGF, and SP. Edema (**A**) was determined by increased paw thickness (mm) in SCI relative to naïve mice. Average concentrations of CGRP (**B**), NGF (**C**), and SP (**D**) found in hindpaw skin for 7 days following SCI. We found that SCI increased paw thickness and concentration of CGRP present in the skin. * = significant difference from naïve, *p*<.05.

Hindpaw edema results from increased release of proinflammatory molecules within the microenvironment of the skin, including CGRP, NGF, and SP [2; 23; 31; 60; 72]. Concentration of each of these respective proteins was measured in hindpaw skin lysate from a cohort of 14 naïve and 12 SCI mice 24 hr following SCI. We chose this timepoint based on our earlier observations that autotomy, spasticity, and edema all develop within this short timeframe. Our analysis did not detect any change in NGF or SP concentrations following SCI. However, CGRP levels in hindpaw skin were significantly elevated following SCI, F(1,25) =4.41, *p*<.05 (**Figures 4B-D**), suggesting the presence of acute-onset CGRP-dependent inflammation within the skin.

### 3.3. Functionally identified afferent populations in naïve and SCI mice 24 hr following injury

Thus far our data suggest a role for peripheral processes in the emergence of spontaneous pain during the acute phase of injury and recovery. To further explore this possibility, we performed *ex vivo* intracellular sharp electrode electrophysiological recordings on 71 cutaneous primary afferents from 19 naïve mice and 81 cutaneous primary afferents from 27 SCI mice. All recordings were performed 24 hr following T9 spinal contusion injury (65 kD, 1s dwell time). We first analyzed afferent response characteristics based, broadly, on responses to specific stimulus modalities. Analyses revealed that 61.19% of naïve neurons responded to temperature, and of that population 58.21% responded to heat and 11.94% responded to cold (**Table 1**). We also found that 88.06% were mechanically sensitive, with 46.27% responding to both mechanical and heat stimulation, 11.94% responding to mechanical and cold stimulation, and 8.96% responding to mechanical, heat, and cold stimulation. Additional analyses demonstrated that 62.40% of SCI neurons were temperature sensitive, with 56.79% responding to heat and 18.52% responding to cold. Furthermore, 83.95% were sensitive to mechanical stimulation, with 44.44% being sensitive to mechanical and heat stimulation, 14.81% responding to mechanical and cold stimulation, and 11.11% were sensitive to mechanical, heat, and cold stimulation, χ^2^=4.08, *p*<.05 (**Table 1**). Statistical comparison of these distributions revealed a significantly greater proportion of SCI afferents that were sensitive to cold stimulation compared to the naïve. No other significant relationships were found.

**Table 1.** Distribution (%) of naïve and SCI afferents that respond to temperature, mechanical, and the combination of temperature and mechanical, stimuli. The percentage of afferents within each category did not differ between conditions except for the percentage of cold responding afferents from SCI mice (bold). *=significantly different from naïve, *p*<.05.

| Stimulus Modality | Naïve % Responders | SCI % Responders |
| --- | --- | --- |
| Temperature | 59.45 | 66.34 |
| Heat | 56.75 | 56.09 |
| Cold | 10.81 | **21.23\*** |
| Mechanical | 85.13 | 82.92 |
| Mechanical + Heat | 41.89 | 42.68 |
| Mechanical + Cold | 10.81 | 14.64 |
| Mecahnical + Heat + Cold | 8.11 | 10.98 |

We further characterized neurons by fiber type (A vs C) and specific stimulus-evoked response using the methods described above to distinguish fiber types (see **Table 2**). The afferent phenotype distribution for naïve neurons included 16.42% A-LTMR, 13.43% A-HTMR, 8.96% CM, 11.94% CH, 0% CC, 2.99% CMC, 37.31% CMH, and 8.96% CMHC neurons. The distribution of SCI afferents included 12.35% A-LTMR, 16.05% A-HTMR, 7.41% CM, 12.35% CH, 3.70% CC, 3.70% CMC, 33.33% CMH, and 11.11% CMHC neurons. Analysis of the distribution of cells between each group revealed a greater percentage of CC neurons present in SCI mice, χ^2^=9.00, *p*<.01. No other comparisons reached statistical significance (**Table 2**).

**Table 2.** Functional population distributions of characterized primary afferent populations from naïve (A) and SCI (B) mice 24 hr following injury. Both groups of mice show similar distributions of A- and C-fibers as well as resting membrane potentials (RMP). However, analysis did reveal an increase in the percentage of CC neurons characterized from SCI mice, *=statistical significance relative to naïve mice, *p*<.05.

| Phenotype | Fiber Type | Function | % Naïve (N=67) | %SCI (N=81) | Naïve RMP (mV) | SCI RMP (mV) |
| --- | --- | --- | --- | --- | --- | --- |
| A-LTMR | A | Low threshold mechanoreceptor | 16.42 | 12.35 | -46.61 ± 3.63 | -56.13 ± 4.20 |
| A-HTMR | A | High threshold mechanoreceptor | 13.43 | 16.05 | -53.45 ± 4.28 | -60.12 ± 3.69 |
| CM | C | Mechanoreceptor | 8.96 | 7.41 | -42.13 ± 6.36 | -50.30 ± 3.41 |
| CC | C | Cold sensitive | 0.00 | **3.70\*** | -- | -39.43 ± 5.43 |
| CH | C | Heat sensitive | 11.94 | 12.35 | -39.65 ± 4.25 | -44.69 ± 4.69 |
| CMC | C | Mechanical + cold sensitive | 2.99 | 3.70 | -42.25 ± 0.67 | -58.50 ± 5.93 |
| CMH | C | Mechanical + heat sensitive | 37.31 | 33.33 | -46.13 ± 1.93 | -44.23 ± 2.02 |
| CMHC | C | Mechanical + heat + cold sensitive | 8.96 | 11.11 | -53.07 ± 3.56 | -49.40 ± 2.47 |

### 3.4. SCI increases mechanical sensitivity in unmyelinated C-fiber nociceptors 24 hr following injury

We assessed mechanical thresholds from 11 naïve and 10 SCI A-LTMR afferents 24 hr following SCI and found no difference in average mechanical thresholds between naïve (4.27 ± 0.47 mN) and SCI (3.67 ± 0.63 mN) neurons (**Figure 5A**). Similar results were also observed when comparing mechanical thresholds of 9 naïve (49.09 ± 17.34 mN) and 13 SCI (33.08 ± 6.58 mN) A-HTMR afferents (**Figure 5B**). Additional analysis of A-LTMR and A-HTMR firing rates in response to mechanical stimulation of the skin at 1, 5, 10, 25, 50, and 100 mN of force revealed no significant differences in firing rates in SCI relative to naïve A-fibers (**Figures 5C, 5D**).

**Figure 5.**
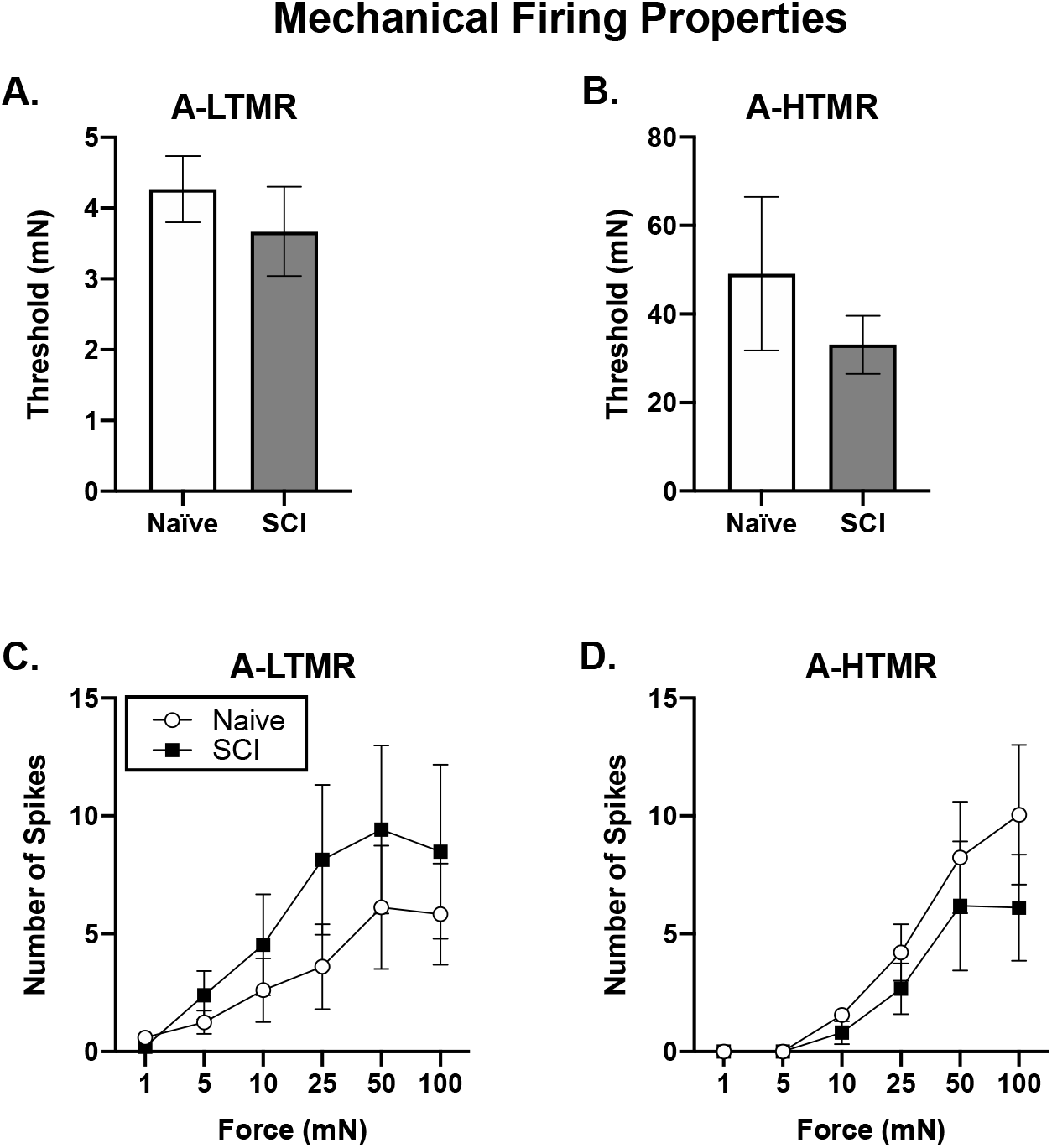
Analysis of mechanically evoked response thresholds and force-dependent firing rates from A-LTMR (**A, C**) and A-HTMR (**B,D**) 24 hr following SCI. No significant differences in average threshold or firing rates emerged, all *p*>.05.

Analysis of average mechanical thresholds from 6 naïve (87.50 ± 11.41 mN) and 6 SCI (75.00 ± 27.54) CM, 2 naïve (62.50 ± 26.52 mN) and 3 SCI CMC (15.00 ± 7.07 mN), 25 naïve (37.60 ± 10.61 mN) and 27 SCI CMH (24.23 ± 5.04 mN), and 6 naïve (36.67 ± 12.78 mN) and 9 SCI CMHC (11.50 ± 5.29 mN) neurons failed to detect significant differences between conditions (**Figures 6A, 6B**). Subsequent analysis of mechanically-evoked firing rates of CMH and CMHC nociceptors showed increased firing in response to 5 and 50 mN of force applied to the skin in SCI CMH neurons and increased responding to 1 and 5 mN of force in SCI CMHC neurons, all Fs>4.86, *p*<.05 (**Figures 6C, 6D**). Despite our ability to obtain mechanical response thresholds from CM and CMC neurons, there was an insufficient number of characterized neurons to perform reliable statistical analysis. Nonetheless, our results suggest that SCI increases mechanical sensitivity in C-fiber nociceptors within 24 hr of injury.

**Figure 6.**
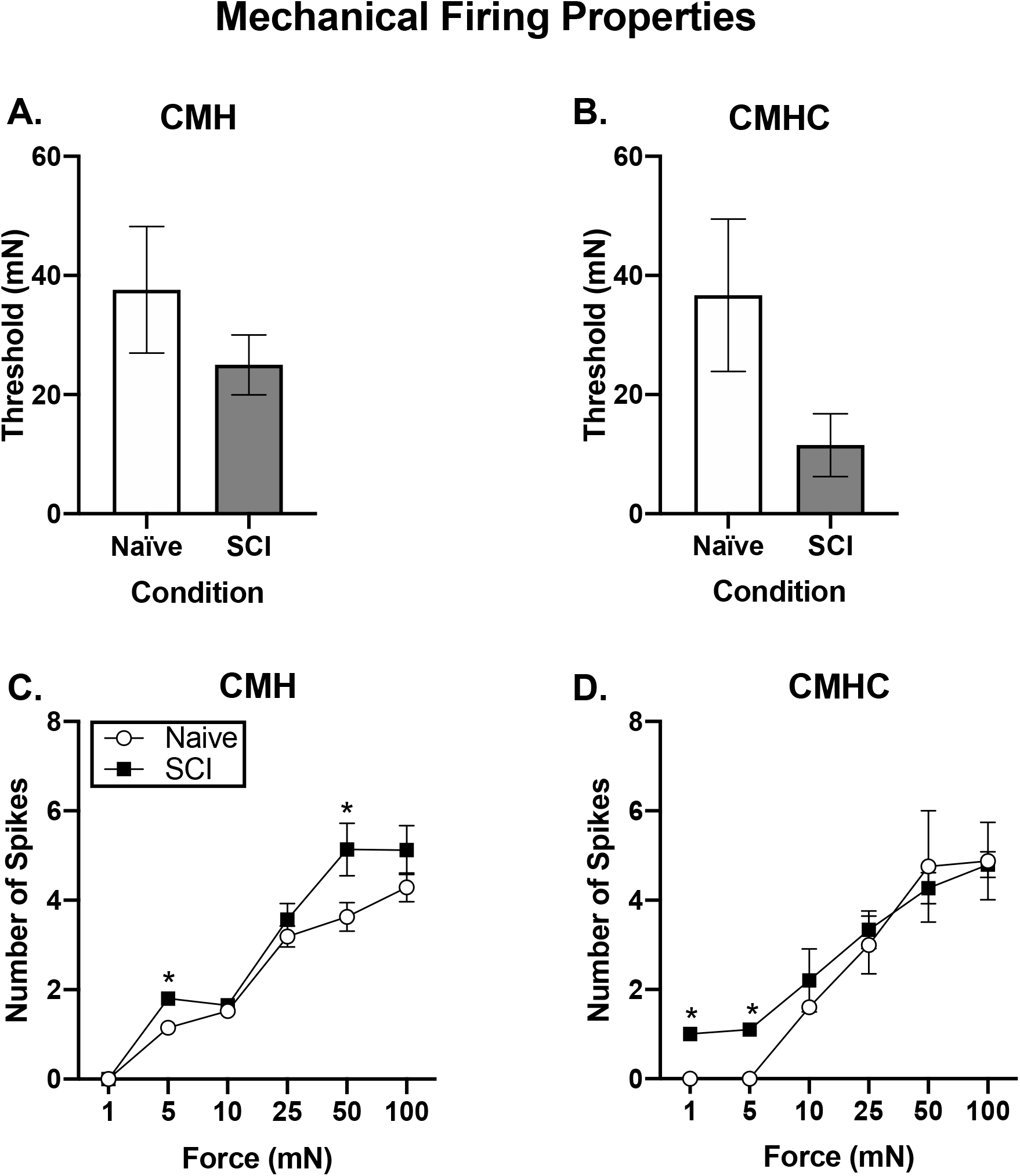
C-fiber mechanical response thresholds and force-dependent firing rates. Response thresholds and firing rates are shown for CMH (**A, C**) and CMHC (**B, D**) nociceptors 24 hr following SCI. While mechanical thresholds were not statistically different between naïve and SCI conditions, we observed a general increase in firing rates as greater force was applied to the skin. We also observed significantly increased firing rates in the lower range of stimulation in SCI relative to naïve neurons. *=significantly different from naïve, *p*<.05.

### 3.5. SCI increases thermal sensitivity in C-fiber nociceptors 24 hr following injury

Temperature thresholds were also examined 24 hr following SCI from 42 naïve and 46 SCI heat sensitive neurons, 8 naïve and 15 SCI cold sensitive neurons, and 4 naïve and 9 SCI neurons that exhibited warming responses. Response to warming was defined as any firing present following an increasing temperature ramp from 0°C to 31°C. We found no differences in response thresholds within heat, cold, or warm temperature-responsive neuronal subtypes. Therefore, we combined afferent subtypes that responded to each stimulus modality and analyzed average heat, cold, and warm thresholds. Analysis of these data failed to detect any significant differences in temperature thresholds between naïve and SCI afferents (**Figure 7A-C**). However, we did find a significantly greater percentage of heat-sensitive SCI nociceptors (21.74%) that responded to warming of the skin following cold stimulation relative to naïve heat-sensitive nociceptors (10.00%), χ^2^=6.25, *p*<.05, with a significantly greater percentage of SCI CMH and CMHC (22.22% and 44.44%, respectively) nociceptors relative to naïve CMH and CMHC (8.00% and 33.33%, respectively) nociceptors, all χ^2^>3.68, *p*<.05. In addition, analysis of thermally-evoked firing rates showed that SCI CMH and CMHC nociceptors exhibited significantly increased firing rates in response to temperatures ranging from 31-38°C and 41-51°C. Additional analysis found that SCI resulted in a significant increase in firing rate in CH nociceptors following 33-35°C and 49-50°C relative to naïve CH nociceptors, all Fs >2.27, *p*<.01 (**Figure 8A-C**), collectively demonstrating that heat sensitivity is increased in thermally responsive nociceptors following SCI.

**Figure 7.**
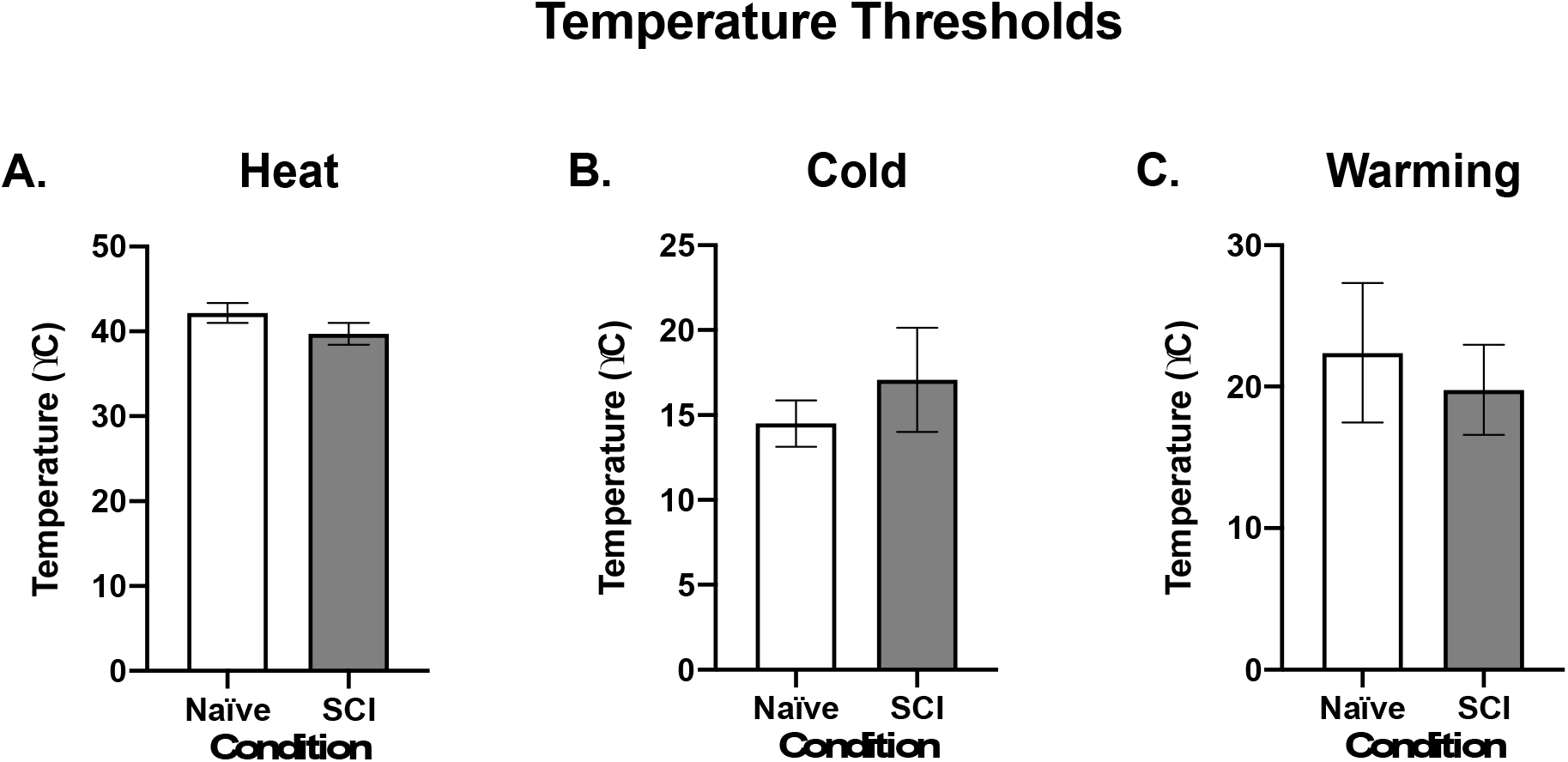
C-fiber temperature response thresholds. Average response thresholds following heat (**A**), cold (**B**), and warming (**C**) stimulation were assessed 24 hr following SCI. No stimulus-specific differences were observed when comparing naïve and SCI neurons, all *p*>.05.

**Figure 8.**
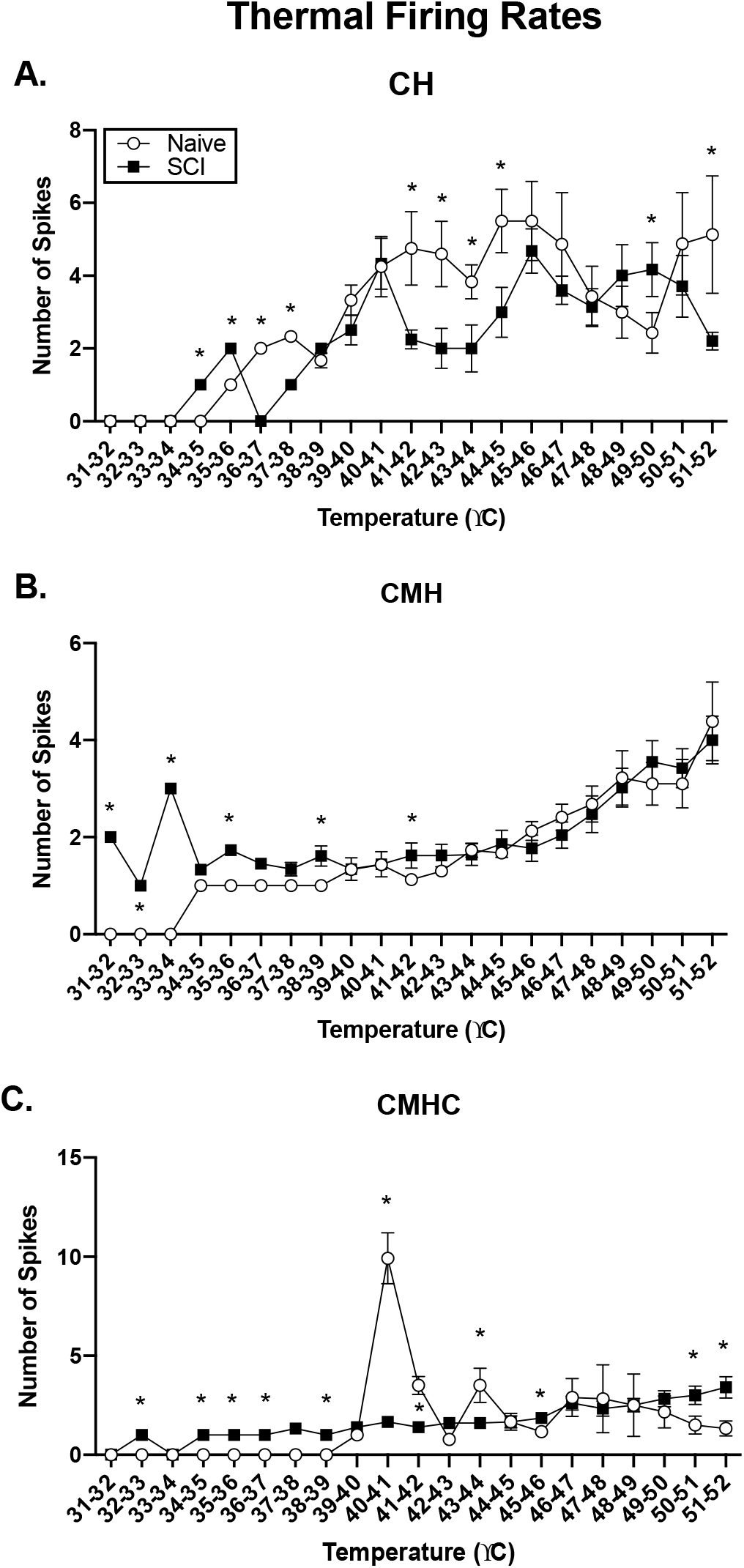
Firing rates for thermally-sensitive nociceptors in response to increasing temperature. Firing rates are depicted for CH (**A**), CMH (**B**), and CMHC (**C**) neurons in response to a 12 s thermal ramp ranging from 31-52°C. In general, we found that SCI increased thermal sensitivity, and SCI neurons fired in response to lower stimulation than naïve nociceptors. *=significantly different from naïve, all *p*<.01.

### 3.6. SCI increases spontaneous firing in C-fiber nociceptors 24 hr following injury

Previous research has shown that SCI increases the incidence of SA and AD in putative nociceptors within days of injury and that SA and AD can persist for several months [8; 9; 58; 86; 89]. The prevalence of AD in nociceptors following SCI is has not been well characterized, and it is not known whether SA and AD are interrelated, whether SA and AD are characteristics of a certain functional population of afferents, or whether an association exists between SCI-induced alterations in thermal and mechanical response thresholds and the incidence of SA or AD. To examine these issues, we assessed the presence of SA and AD in all electrophysiologically characterized neurons and found a significantly greater percentage of SCI neurons (39.02%) exhibited SA relative to naïve neurons (1.35%), χ^2^=684.5, *p*<.0001. When we analyzed the presence of SA in specific functional subtypes of afferents, we found that 0/11 naïve and 2/10 SCI LTMRs, 0/9 naïve and 0/13 SCI HTMRs, 1/6 naïve and 5/6 SCI CMs, 0/11 naïve and 6/10 SCI CHs, 0/2 naïve and 1/3 SCI CMCs, 0/25 naïve and 14/27 SCI CMHs, and 0/6 naïve and 4/9 SCI CMHCs exhibited SA. Analysis of these data revealed a significant increase in the occurrence of SA in SCI LTMR, CM, CH, CMH, and CMHC neurons relative to the same functional populations of naïve neurons, all χ^2^>4.00, *p*<.05 (**Figure 9A**). Analysis of the prevalence of AD revealed similar results, demonstrating that while AD was absent in all naïve neurons, we did observe AD in 41.46% of SCI neurons. Quantification of the presence of AD within in each functional subclass of SCI neurons showed that 1/10 LTMRs, 0/13 HTMRs, 4/6 CMs, 0/10 CHs, 1/3 CMCs, 23/27 CMHs, and 5/9 CMHCs exhibited AD (**Figure 9B**). Analysis revealed statistically significant increases in AD occurrence in SCI CM, CMH, and CMHC relative to naïve neurons in the same functional categories.

**Figure 9.**
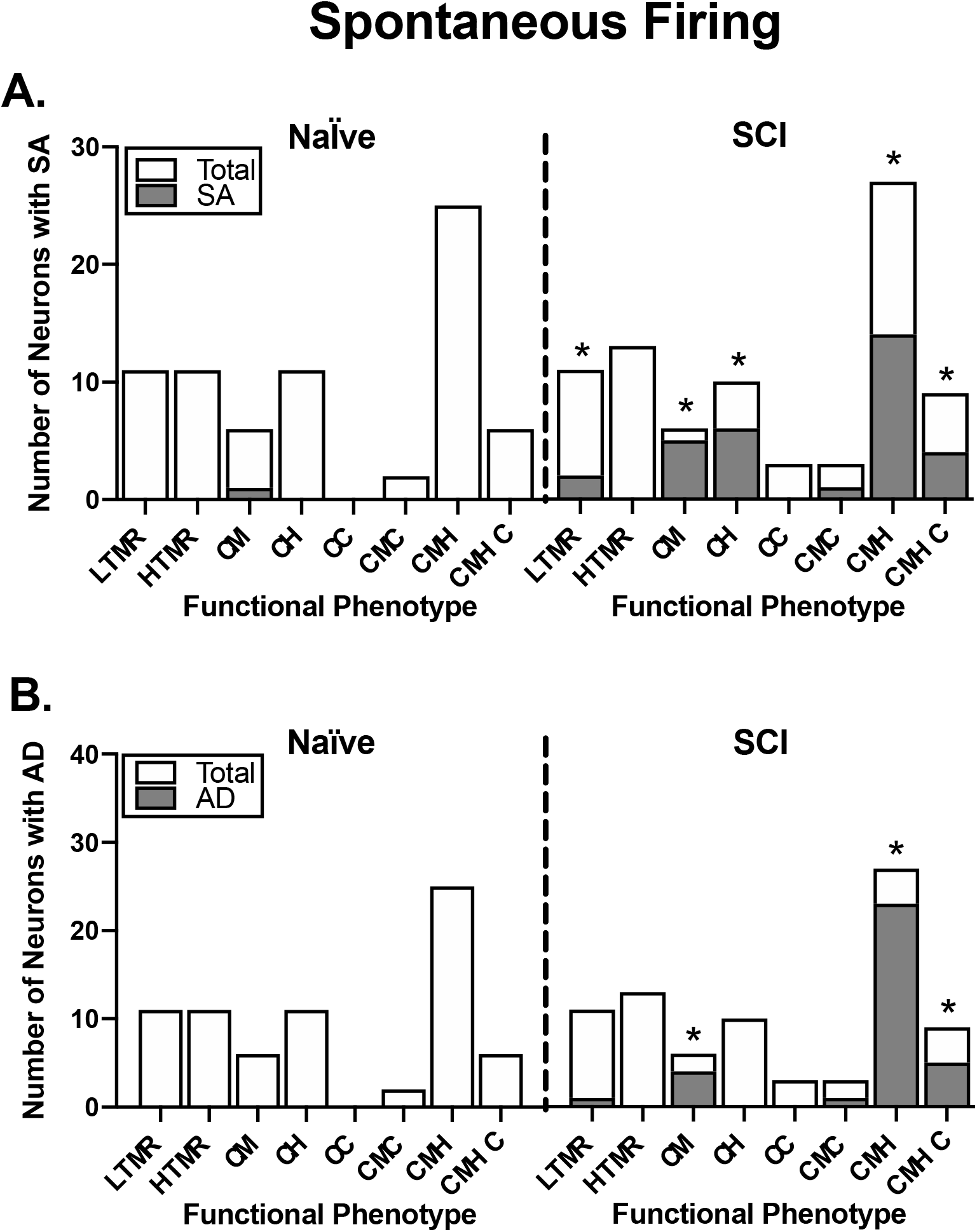
Spontaneous activity (SA) and afterdischarge (AD) in functional subtypes of afferents. **A**) incidence of SA in characterized naïve and SCI afferents. Data are depicted as number of neurons that exhibited SA (gray bars) relative to total of characterized neurons (white bars). **B**) incidence of AD in characterized naïve and SCI neurons depicted as number of neurons that exhibited AD (gray bars) relative to total of characterized neurons (white bars).

It is possible that the increase in SA and AD following SCI was due to alterations in the intrinsic electrophysiological properties of neurons, potentially resulting in a relative state of depolarization and a subsequent decrease in firing threshold. To examine this possibility, we compared resting membrane potentials (RMPs) in each neuronal subtype across injury conditions. We found no significant differences in RMPs in any subpopulation of afferents following SCI compared to naïve subjects (**Table 2**).

These above results demonstrate that SA and AD occur in similar functional distributions of nociceptive afferents in response to SCI. Pearson correlations were then performed to determine whether there was an association between the presence of SA and AD in a given cell and failed to detect any significant relationships (r_P_=0.048, *p*>.05). We then examined whether nociceptor phenotype, mechanical sensitivity, or heat sensitivity was significantly correlated with SA or AD using linear regression. We failed to resolve any significant association between specific nociceptor subtype or mechanical sensitivity with the incidence of SA or AD (all r_P_ <0.32, *p*<.05). However, our analysis did reveal a significant inverse relationship between heat sensitivity and the presence of SA (r_P_ = −0.290, *p*<.05) and significant positive relationship between heat sensitivity and the presence of AD (r_P_ = 0.300, *p*<.05). Collectively, these results suggest that SA and AD are unrelated to each other and that a population of low threshold heat-responsive nociceptors are more likely to exhibit SA, while a separate population of high threshold heat-responsive nociceptors are more likely to exhibit AD.

## 4. Discussion

### 4.1. SCI increases spontaneous pain behavior, neurogenic-like inflammation, and nociceptor sensitivity

Chronic pain occurs at an alarmingly greater percentage in the spinally injured relative to the general chronic pain population [16; 21; 29; 39; 62; 71], is often poorly managed, and is associated with poor prognoses for recovery [5-7; 34; 37]. The mechanisms underlying SCI-induced pain are initiated rapidly following injury [9], and our present results demonstrate that autotomy, spasticity, hindpaw edema, and increased CGRP levels occurred 1-7 days following injury, suggesting that injured animals experience ongoing pain via a neurogenic inflammation-like process that could augment nociceptor response properties.

Given the above results we next examined potential SCI-induced alterations in nociceptor response properties during the first 24 hr following SCI. The majority of characterized neurons from both naïve and SCI mice were mechanically sensitive, with a smaller, but significant, proportion of afferents that were temperature sensitive. The distribution of temperature-responsive neurons was similar between conditions, with the exception of increased cold-sensitive afferents after SCI. However, this finding should be interpreted with some caution because we did not characterize any CC neurons from naïve mice, which is unusual.

Alterations in afferent response characteristics appear to emerge due to increased sensitivity within specific functional populations of neurons. We did not observe changes in mechanical response thresholds or firing rates in A-fiber afferents. We did observe increased action potential (AP) firing in response to mechanical force applied to the skin that was independent of changes in average mechanical response thresholds in C-fiber afferents, with CMH and CMHC nociceptors exhibiting a leftward shift in mechanical sensitivity; CMH nociceptors exhibited increased firing in response to 5 mN of force and CMHC nociceptors exhibited increased firing in response to 1 and 5 mN of force applied to the skin. Importantly, these increases in firing rates occurred at or near innocuous forces of stimulation, suggesting that normally innocuous mechanical stimulation may heighten nociceptor responding that potentially contributes to increased pain (e.g., allodynia) after SCI. SCI also increased AP firing in response to 50 mN of applied force in CMH nociceptors, indicating that activation of this specific subpopulation of nociceptors by noxious mechanical stimulation increases activity that may promote development of hyperalgesia.

Application of hot, warm, and cold stimuli to the skin did not reveal any significant differences in average response thresholds between naïve and SCI mice. However, we did find an increase in the number of nociceptors that responded to warming of the skin following SCI, suggesting an overall increase in thermal sensitivity. We also found that thermally sensitive nociceptor populations (e.g., CH, CMH, and CMHC) exhibited leftward shifts in their respective temperature response curves following SCI; SCI CH nociceptors exhibited an increase in firing rate at 34°C, while naïve CHs began firing at 35°C; SCI CMH nociceptors began firing at 31°C, while naïve CMHs began firing at 34°C; and SCI CMHC nociceptors showed modest, but significant, increases in firing at 32°C, while naïve CMHCs began firing at 39°C. Notably, exposure to increasing temperatures produced complex stimulation-response curves that could result from system-wide adaptations in response to injury, including spinal shock or energy deficiencies related to increased demands during the healing process and SCI-induced mitochondrial dysfunction.

### 4.2. SCI increases spontaneous activity and afterdischarge in specific populations of C-fiber nociceptors

SA is observed in dissociated afferent cell bodies within 3 days of SCI, that persists for at least 8 months, and leads to the development of chronic SCI pain [8; 9; 89]. These findings have been instrumental in establishing a role for peripheral neurons in the development and maintenance of chronic SCI pain. The results detailed here build upon those findings by showing that SA and AD emerge within the first 24 hr of injury in CM, CH, CMH, CMHC, and to a lesser extent, A-LTMR neurons. We also show that AD is observed in CM, CMH, and CMHC neurons. While SA and AD are found in similar functional populations of neurons, their relative distributions are not identical. The overall presence of SA and AD are not correlated with one another, suggesting that independent molecular processes may be responsible for their development, even within the same cell. In addition, we found independent relationships between SA, AD, and thermal, but not mechanical, sensitivity, where SA was correlated with lower heat thresholds while the presence of AD was correlated with higher heat thresholds. While further work is needed to detail these mechanisms, our results indicate that low threshold nociceptors may fire continuously in the presence and absence of stimulation, while high threshold nociceptors may continue to fire despite termination of a heat stimulus, both of which may initiate miscoding of sensory input. Moreover, because below-level nociceptors exhibit increases in excitability, it is also possible that SCI causes widespread thermal and mechanical sensitivity between systems (e.g., musculoskeletal, cutaneous, visceral) that may lead to indiscriminable, widespread pain.

### 4.3. Potential mechanisms responsible for central recruitment of nociceptive afferents and subsequent sensitization

The development of SA results from depolarizing RMPs, reductions in AP threshold, and the occurrence of depolarizing spontaneous fluctuations (DSFs) [56]. We did not observe any obvious DSFs or alterations in RMPs, which may have resulted from the presence of an intact peripheral sensory circuit that that regulates nociceptor excitability via interactions at sites of peripheral and central innervation. Supporting this, prior work has shown that SCI increases the early release of proinflammatory cytokines, such as IL-1β, in the spinal cord that increase afferent sensitivity [55], and we show increased CGRP expression, which recruits inflammatory/immune cells that release other neuroactive molecules, such as prostaglandins, that can further increase nociceptor excitability [12; 20; 43; 66]. Moreover, central and peripheral inflammatory stimulation may drive ectopic afferent activity that increases nociceptor sensitivity and excitability similar to what has been observed following induction of wind-up, central sensitization, and long-term potentiation [19; 40; 41; 51; 67-69; 73; 76; 80; 81; 85; 87; 88].

### 4.4. Functional and clinical importance of altered evoked and spontaneous nociceptor firing properties

Activation of nociceptors following SCI may create a “reverberatory loop”, where antidromic nociceptor APs generated in the injured spinal cord causes release of neuroactive molecules from peripheral terminals (e.g., CGRP) into their targets of innervation [82], causing generation and propagation of orthodromic nociceptor APs into the spinal cord that fosters the development and/or maintenance of an excitatory environment within the spinal cord. Consequently, breaking this self-perpetuating loop of nociceptive communication could serve as a valuable mechanistic target for treating chronic SCI pain by silencing nociceptor output. Furthermore, treatments that disrupt nociceptor could be administered during the first few hours following SCI, when nociceptor output is increasing, with the intent of preventing pain development. In laboratory settings this approach has been highly successful and experimental work has repeatedly shown that prevention of chronic pain development is possible while reversal of chronic pain is incredibly difficult. Consequently, SCI is a unique injury scenario where intervention can be administered prophylactically, close to the time of injury.

Activation of nociceptors by innocuous mechanical or thermal stimuli suggests that numerous endogenous or environmental stimuli have the potential to cause pain, including normal body temperature, limb movement, mechanical forces involved in gastrointestinal function, contact with wheelchairs, bedsheets, clothing, and administration of necessary medical procedures (e.g., catheterization, IV placement, etc.) could have detrimental effects for patient pain burden and recovery. Consequently, therapeutics designed to quiet nociceptor activity during the acute phase of injury may have broad reaching effects by improving patient recovery prognosis and quality of life.

## Acknowledgments

The authors have no conflicts of interest to report.

Funding for the current studies was provided by NINDS R03NS096454, NINDS R21NS104789, NIGMS P20GM103418, NIH U54 HD 090216, Rita Allen Foundation Award in Pain, KUMC Biomedical Research Training Program, and the Madison and Lila Self Graduate Fellowship.

The authors would like to thank Ms. Leena Kader, Ms. Taylor Messer, and Mr. Adam Willits for their helpful comments on previous versions of this manuscript.

